# IRONMAN Tunes Responses to Iron Deficiency in Concert with Environmental pH

**DOI:** 10.1101/2021.02.16.431461

**Authors:** Chandan Kumar Gautam, Huei-Hsuan Tsai, Wolfgang Schmidt

**Affiliations:** Molecular and Biological Agricultural Sciences Program, Taiwan International Graduate Program, Academia Sinica and National Chung-Hsing University, Taipei 11529, Taiwan; Graduate Institute of Biotechnology, National Chung-Hsing University, Taichung 40227, Taiwan; Institute of Plant and Microbial Biology, Academia Sinica, Taipei 11529, Taiwan; Biotechnology Center, National Chung-Hsing University, Taichung 40227, Taiwan; Genome and Systems Biology Degree Program, College of Life Science, National Taiwan University, Taipei 10617, Taiwan

## Abstract

Iron (Fe) is an essential mineral element which governs the composition of natural plant communities and limits crop yield in agricultural ecosystems due to its extremely low availability in most soils, particularly at alkaline pH. To extract sufficient Fe from the soil under such conditions, some plants including *Arabidopsis thaliana* secrete Fe-mobilizing phenylpropanoids, which mobilize sparingly soluble Fe hydroxides by reduction and chelation. We show here that ectopic expression of the *IRONMAN* peptides *IMA1* and *IMA2* improves growth on calcareous soil by inducing the biosynthesis and secretion of the catecholic coumarin fraxetin (7,8-dihydroxy-6-methoxycoumarin) through increased expression of *MYB72* and *SCOPOLETIN 8-HYDROXYLASE* (*S8H*), a response which is strictly dependent on elevated environmental pH (pH_e_). By contrast, transcription of the cytochrome P450 family protein *CYP82C4,* catalyzing the subsequent hydroxylation of fraxetin to sideretin, which forms less stable complexes with iron, was strongly repressed under such conditions. Luciferase reporter assays in transiently transformed protoplasts showed that IMA1/IMA2 peptides are translated and modulate the expression of *CYP82C4* and *MYB72* by acting as transcriptional coactivators. It is concluded that IMA peptides regulate processes supporting Fe uptake at both acidic and elevated pH by controlling gene expression upstream of or in concert with a putative pH_e_ signal to adapt the plant to the prevailing edaphic conditions. This regulatory pattern confers tolerance to calcareous soils by extending the pH range in which Fe can be efficiently absorbed from the soil. Altering the expression of IMA peptides provides a novel route for generating plants adapted to calcareous soils.

**One sentence summary:** Ectopic expression of IRONMAN peptides improves growth under iron-limiting conditions by inducing responses to limited iron availability in accordance with the environmental pH.

The author responsible for distribution of materials integral to the findings presented in this article in accordance with the policy described in the Instructions for Authors (www.plantphysiol.org) is: Wolfgang Schmidt.

## INTRODUCTION

In natural ecosystems, growth on calcareous, i.e. carbonate-rich, alkaline soils, is restricted to so-called calcicole (chalk-loving) plants, while calcifuge (chalk-fleeing) species are excluded from such habitats (Lee, 1998; Grime and Hodgson, 1969). Among other factors, the ability to thrive on calcareous substrates has been attributed to a particularly efficient strategy to extract iron (Fe) from the soil, an essential mineral nutrient with extremely limited solubility in most conditions. In aerated soils, alkalinity decreases the availability of Fe to levels that are several orders of magnitude below the requirement of the plants (Lindsay and Schwab, 1983; Tyler, 1996). Species that are not well adapted to calcareous soils, as for example crops such as fruit trees and soybean or taxa which originate from acidic habitats, develop lime-induced chlorosis caused by insufficient uptake, maldistribution, or immobilization of Fe (Zohlen and Tyler, 2003).

The underlying causes for the superior Fe acquisition efficiency of calcicole species still remain largely enigmatic. Generally, gramineous species (Poaceae) have been regarded as being more Fe-efficient under alkaline condition than non-grasses, a distinction that has been associated with the mechanism by which Fe is mobilized by grass roots. In contrast to all other land plants, grasses have adopted a chelation-based Fe uptake system that relies on the synthesis and secretion of Fe-chelating phytosiderophores, a mechanism that has been designated as strategy II (Römheld and Marschner, 1986). Phytosiderophores are mugineic acid derivates that form soluble complexes with ferric Fe, which are fairly stable over a wide pH range (Shi et al., 1988). Conversely, non-grasses have adopted an Fe acquisition strategy which depends on the orchestrated action of enzymatic reduction of ferric chelates, H^+^-ATPase-mediated acidification of the rhizosphere, and secretion of low-molecular-weight Fe-mobilizing compounds such as flavins or phenylpropanoids, a system that is referred to as strategy I (Römheld and Marschner, 1986). In Arabidopsis, chelated ferric Fe is reduced by the plasma membrane-bound FERRIC REDUCTION OXIDASE2 (FRO2; Robinson et al., 1999) and the liberated Fe^2+^ is subsequently taken up from the soil solution by IRON-REGULATED TRANSPORTER1 (IRT1; Eide et al; 1996; Vert et al., 2002). The FRO2-mediated reduction of ferric Fe is supported by proton extrusion via the P-type H^+^-ATPase AHA2 (Santi and Schmidt, 2009), a process that creates a slightly acidic pH milieu in the apoplast that is favorable for the uptake of Fe. FRO2-mediated ferric reduction is compromised at circumneutral or alkaline pH ranges (Susín et al., 1996; Lucena et al., 2007; Waters et al., 2018), rendering the strategy I system ineffective under such conditions.

In Arabidopsis, the genes mediating the Fe deficiency responses are under the control of a heterodimer consisting of the master transcription factor FER-LIKE IRON DEFICIENCY INDUCED TRANSCRIPTION FACTOR (FIT; bHLH29) and one out of four clade Ib bHLH proteins (bHLH38, bHLH39, bHLH100, and bHLH101) (Colangelo and Guerinot, 2004; Yuan et al., 2008; Sivitz et al., 2012; Wang et al., 2013). The expression of FIT, in turn, and that of all Ib bHLH proteins, is controlled by UPSTREAM REGULATOR OF IRT1 (URI/bHLH121) in concert with IAA-LEUCINE RESISTANT3 (ILR3/bHLH105) (Kim et al., 2019; Gao et al., 2020a; Lei et al., 2020). FIT is negatively regulated by the clade IVa bHLH transcription factors bHLH18, bHLH19, bHLH20, and bHLH25 (Cui et al., 2018), and by the E3 ligases BTSL1 and BTSL2 (Rodríguez-Celma et al., 2019), which target FIT to degradation via the 26S proteasome. A novel family of peptides designated IRONMAN/FE-UPTAKE-INDUCING PEPTIDE (IMA/FEP), control Fe uptake presumably via regulation of clade Ib bHLH proteins (Grillet et al., 2018; Hirayama et al., 2018). IMA/FEP peptides are ubiquitous across land plants and share a highly conserved consensus motif at the C-terminus, which is necessary and sufficient for IMA function. The expression of *IMA1* and *IMA2* is directly controlled by bHLH121 (Gao et al., 2020a). Ectopic expression of *IMA1* overrides the repression of Fe uptake by sufficient internal Fe levels, induces ferric reduction activity, and improves growth on media with low Fe solubility (Grillet et al., 2018). High ferric reduction activity is, however, insufficient to confer calcicole behavior (Schmidt and Fühner, 1998; Terés et al., 2019), indicating that other mechanisms are employed to enable plants to thrive under such conditions. Recent studies have shown that growth of non-grasses on calcareous soils requires the synthesis and secretion of species-specific, low-molecular-weight compounds, which increase the solubility of ferric hydroxides by reduction and chelation and greatly extend the pH range in which Fe can be efficiently mobilized (reviewed by Chen et al, 2017; Tsai and Schmidt, 2017; Robe et al., 2020b). In Arabidopsis, Fe deficiency triggers a massive increase in the abundance of enzymes of the phenylpropanoid pathway and an accumulation of the hydroxycoumarin scopoletin (Lan et al., 2011). Scopoletin, however, does not possess Fe-mobilizing properties, which suggests that other compounds derived from this pathway are critical for the acquisition of Fe.

The Fe(II)- and 2-oxoglutarate-dependent dioxygenase FERULOYL-COA 6’-HYDROXYLASE1 (F6’H1) catalyzes the *or*ŵ*o*-hydroxylation of feruloyl-CoA into 6-hydroxyferuloyl-CoA, the first committed step in the biosynthesis of catecholic coumarins derived from scopoletin, and was identified in a genetic screen as being essential for survival on alkaline soil (Schmid et al., 2013). In the light, 6-hydroxyferuloyl-CoA is converted into scopoletin, a reaction that occurs partially spontaneous. In light-protected organs such as roots, scopoletin biosynthesis is dependent on enzymatic catalysis via COUMARIN SYNTHASE (COSY) (Vanholme et al., 2019). Similar to F6’H1, COSY activity is indispensable for growth on alkaline substrates (Vanholme et al., 2019), suggesting that compounds which are produced downstream of scopoletin are required for the mobilization of Fe. Hydroxylation of scopoletin mediated by another Fe(II)- and 2-oxoglutarate-dependent dioxygenase, SCOPOLETIN 8-HYDROXYLASE (S8H), yields fraxetin (7,8-dihydroxy-6-methoxycoumarin), a potent Fe chelator (Siwinska et al., 2018; Tsai et al., 2018, Rajanik et al., 2018). A further hydroxylation step, mediated by CYTOCHROME P450 FAMILY 82 SUBFAMILY C POLYPEPTIDE4 (CYP82C4), converts fraxetin into a novel catecholic coumarin referred to as sideretin (5,7,8-trihydroxy-6-methoxycoumarin) (Rajanik et al., 2018). Secretion of scopoletin, fraxetin, and sideretin occurs via PLEIOTROPIC DRUG RESISTANCE9 (PDR9) (Fourcroy et al., 2014; Ziegler et al., 2017; Rajniak et al., 2018).

Fraxetin and sideretin feature adjacent hydroxyl groups which bind Fe efficiently and constitute the major Fe-mobilizing coumarins derived from this pathway (Schmid et al., 2014; Rajanik et al., 2018; Tsai et al., 2018; Robe et al., 2020b). In particular, fraxetin secretion appears to be important for plant edaphic adaptation. Growth and chlorophyll content correlated with fraxetin secretion across a suite of Arabidopsis accessions grown on high pH media with restricted Fe availability, indicating that this process is critical for the ability to thrive on alkaline soils (Tsai et al., 2018). Correspondingly, the better Fe uptake of Arabidopsis demes with high carbonate content in their native habitat relative to demes native to silicious soils was found to be due to higher proton extrusion and higher fraxetin secretion (Terés et al., 2019). Supplementing the growth media with fraxetin or growth of sensitive plants from silicious soils alongside carbonate-adapted plants enhanced Fe and chlorophyll levels of the sensitive plants, underscoring a key role for Fe-mobilizing coumarins in the mobilization of Fe from recalcitrant pools (Terés et al., 2019).

Here, we set out to explore the basis on which IMA is conferring calcicole behavior to *Arabidopsis thaliana,* a calcicole species. We demonstrate that overexpression of either *IMA1* or *IMA2* is sufficient to activate all Fe deficiency responses except for fraxetin secretion, which is modulated by the environmental hydrogen activity (pH_e_). Elevated pH_e_ promotes the expression of *S8H* and represses the transcription of *CYP82C2,* thereby activating a separate, calcicole branch of the strategy I-type Fe deficiency response.

## RESULTS

### IMA1 and IMA2 are Closely Co-Expressed Paralogs

IMA1 and IMA2 are highly similar peptides harboring a conserved C terminus that has been associated with Fe uptake across land plants (Grillet et al., 2018). The two peptides share 82% sequence identity and are robustly induced by Fe deficiency in both roots and shoots (Fig. 1A; Rodríguez-Celma et al., 2013a; 2013b). Publicly available RNA-seq data reveal similar organ- and tissue-level expression of both genes (Supplementary Fig. S1). To infer organ-specific coexpression networks around *IMA1* and *IMA2,* we downloaded and normalized a total of 5,556 RNA-seq data sets from the Sequence Read Archive (SRA) hosted by the National Center for Biotechnology Information (NCBI). Networks were constructed with the MACCU toolbox (Lin et al., 2011) based on pair-wise comparison of the co-expression relationships of Arabidopsis genes expressed in either roots or leaves with a Pearson coefficient > 0.7 using tissue-specific data sets (1,194 for roots and 641 for leaves). In both roots and leaves, *IMA1* and *IMA2* are coexpressed with a suite of other putative or validated regulators of cellular Fe homeostasis such as *IMA4*, *bHLH38*, *bHLH39,* and *IRON RESPONSIVE PROTEIN3* (*IRP3;*At2g14247) (Fig. 1B). Together with IMAs, IRPs were identified in transcriptional surveys as being highly Fe responsive in both roots and shoots (Rodríguez-Celma et al., 2016a). IRP1 and IRP2 have been renamed to IMA1 and IMA2 after elucidation of their function (Grillet et al., 2018); the biological role of IRP3-IRP6 remains to be elucidated. In roots, *IMA1/IMA2* are further coexpressed with *NICOTIANAMINE SYNTHASE4 (NAS4)* and *IRP6* (At5g05250). In leaves, the expression of *IMA1/IMA2* is associated with *IMA3* and *bHLH100* (Fig. 1B).

**Figure 1.**
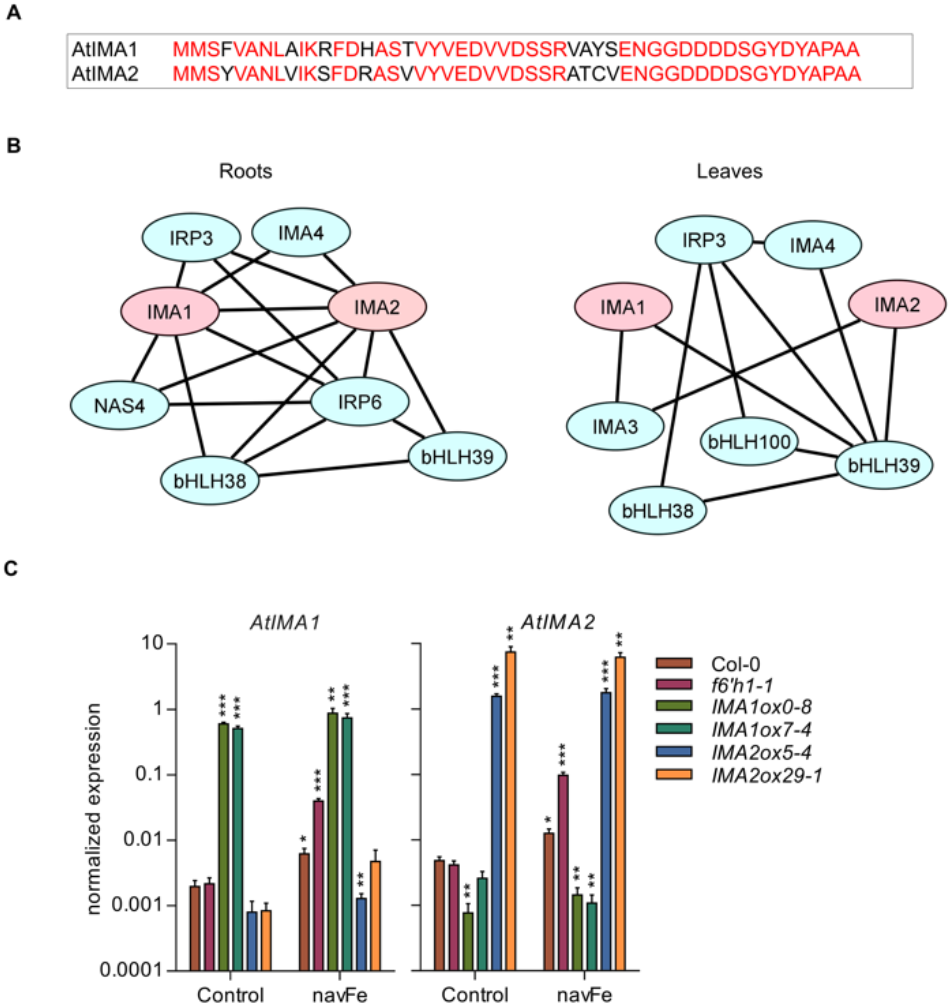
Sequences, co-expression networks, and expression levels of *IMA1* and *IMA2* in transgenic plants. A, Alignment of IMA1 and IMA2 peptides. B, Co-expression network around *IMA1* and *IMA2* in roots and leaves. Networks were constructed with the MACCU toolbox (Lin et al., 2011) against customized organ-specific data bases. Genes with a Pearson coefficient of > 0.7 for pairwise expression were included in the network. C, Expression of *IMA1* and *IMA2* in the wild type, *f6’h1* mutant plants, and transgenic lines overexpressing either *IMA1 (IMA1ox}* or *IMA2 (IMA2ox).* Expression was measured in roots of 12-d-old seedling grown on ES or navFe media for 3 days after 9 days of pre-cultivation on ES media. Each bar represents the mean ± SE of four independent experiments. Asterisks indicate significant differences from the wild type in each treatment: *, *P* ≤ 0.05; **, *P* ≤ 0.01; ***, *P* ≤ 0.001.

For further studies on the function of IMA1 and IMA2, we generated transgenic lines that expressed either gene under the control of the CaMV 35S promoter, leading to an increase in *IMA1* transcript levels between ca. 260-to 300-fold in *IMA1* overexpression lines *(IMA1ox)* and 320-to 1,500-fold of *IMA2* in *IMA2ox* lines compared to the wild type (Fig. 1C). In wild-type plants, growth on media with restricted Fe availability (navFe media; 10 μM FeCl_3_, pH 7.0) increased *IMA1* and *IMA2* expression by ca. 3-fold; no further increase in IMA expression was observed in *IMA1ox* lines. Mutants harboring a defect in the expression of *F6’H1* showed a more than 20-fold increased transcript levels of both *IMA* genes relative to wild-type plants when grown under navFe conditions, possibly reflecting a severely low Fe status of the mutant plants (Fig. 1C).

### Overexpression of *IMA1/IMA2* Supports Fe Uptake at Elevated pH

Growing plants on soil supplemented with calcium oxide (CaO) resulted in pronounced growth reduction in wild-type plants, which was more severe at higher (1%) CaO levels (Fig. 2A). Plants grown on alkaline soil produced chlorotic leaves, indicative of restricted Fe availability. Mutants defective in the expression of *F6’H1* showed a more severe phenotype than the wild type, indicating that secretion of coumarins is critical for survival under such conditions. *IMA1ox* and *IMA2ox* lines showed markedly improved growth when compared to wild-type plants, a discrepancy which was more obvious in soil supplemented with 1% CaO (Fig. 2A). *IMAox* plants grown under such conditions were bigger and greener than the heavily chlorotic wild-type plants, an observation which is likely attributable to improved Fe acquisition.

**Figure 2.**
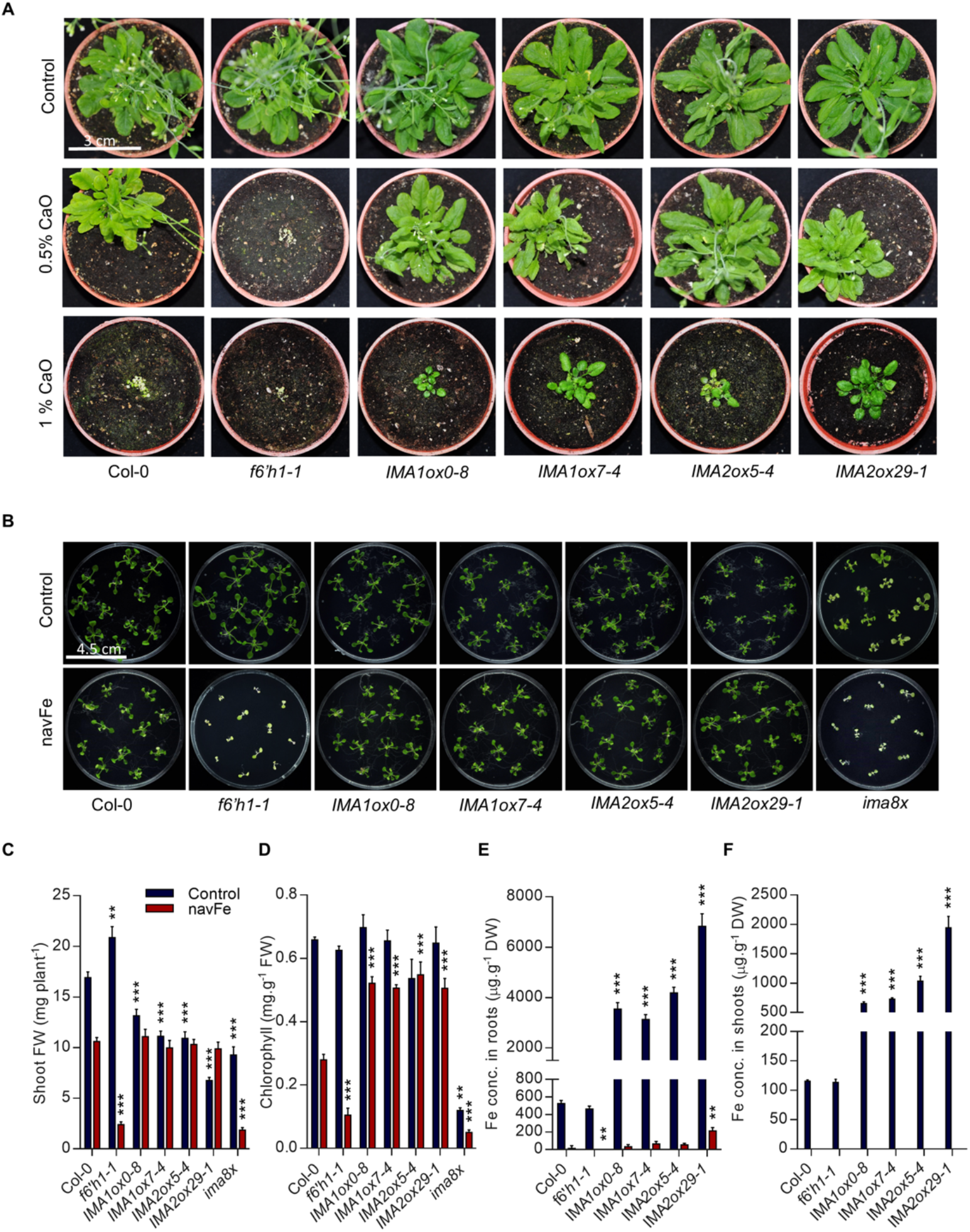
Overexpression of *IMA1* and *IMA2* confers tolerance to alkaline conditions. A, Phenotypes of plants grown for 8 weeks on alkaline soil supplemented with CaO. B, Phenotypes of plants grown on ES (50 μM FeEDTA, pH 5.5) or navFe (10 μM FeCl_3_, pH 7.0) media for 2 weeks. C, Shoot fresh weight. D, Chlorophyll concentration. E, F, Iron concentration in roots (E) and shoots (F). Each bar represents the mean ± SE of four (seven for shoot fresh weight) independent experiments. Asterisks indicate significant differences from the wild type in each treatment: *, *P* ≤ 0.05; **, *P* ≤ 0.01; ***, P ≤ 0.001.

Growing plants on navFe media reduced growth of the wild type by about 40% (Fig. 2B,C). When grown on media supplemented with 50 μM FeEDTA at pH 5.5, *f6’h1* mutant plants were significantly bigger than the wild type, matching previous findings (Rodríguez-Celma et al., 2013b). Growth of *f6’h1* mutants on navFe media was, however, massively impaired, leading to a ca. 90% decrease in shoot fresh weight at the end of the experimental period (Fig. 2B,C). *IMA1/IMA2ox* lines grew slower than the wild type under Fe-sufficient conditions (ca. 40% reduction in shoot fresh weight on average), but did not show any significant growth reduction on navFe media. Octuple *ima8x* mutants, defective in the expression of all eight *IMA* genes, were highly chlorotic under all conditions, a phenotype which was more severe when grown on navFe media where *ima8x* resembled*f6’h1* mutant plants (Fig. 2B,C). Growth on navFe media further caused a pronounced decrease in chlorophyll concentration in wild-type and, more dramatic, in *f6’h1* mutant plants. *IMA1*/*IMA2ox* lines exhibited a much less severe reduction in chlorophyll levels. On average, chlorophyll concentration of *IMAox* lines was decreased by 17% vs. 58% in wild-type and 83% in *f6’h1* mutant plants (Fig. 2D). The lowest chlorophyll concentration was observed in *ima8x* mutants (Fig. 2D).

Histochemical detection of Fe by Perls staining revealed massive accumulation of Fe in roots and shoots of both *IMA1ox* and *IMA2ox* lines under Fe-sufficient conditions and when grown on media supplemented with excess Fe (400 μM FeEDTA), suggesting that both genes play at least partly redundant roles in the regulation of Fe uptake (Supplementary Fig. S2). Growing plants on navFe media abolished Fe staining completely in both wild-type and *f6’h1* mutant plants. In *IMA1/IMA2ox* lines, staining was weak but still above the detection limit (Supplementary Fig. S2), indicating that under such conditions Fe uptake is less restricted in lines overexpressing either *IMA1* or *IMA2.* Quantifying Fe levels in the genotypes under investigation supported the semi-quantitative analysis and revealed a massive increase in root and shoot Fe concentration in *IMA1/IMA2ox* lines under Fe-sufficient conditions relative to wildtype plants (Fig. 2E,F). When grown on navFe media, Fe was barely (in roots) or not at all (in shoots) detectable (Fig. 2E,F). To quantify Fe levels under these conditions, we analyzed plants that were pre-grown on Fe-sufficient media for 7 days and subsequently transferred to navFe media for additional five days. Iron levels remained low in all genotypes under these conditions, but *IMA1/IMA2ox* lines exhibited an on average more than 30-fold increase in Fe levels in roots and in shoots, suggesting that IMA peptides particularly support Fe uptake under conditions of restricted Fe availability (Supplementary Fig. S3).

### IMA Regulates Fe Acquisition at Different Levels

To investigate the effect of IMA1 and IMA2 on the different Fe acquisition processes, we first analyzed the capacity of the lines under study to acidify the rhizosphere. Under Fe-sufficient conditions, a low constitutive acidification confined to the root tip was recorded for wild-type and *f6’h1* mutant plants; roots of *f6’h1* plants were slightly more active (Fig. 3A). *IMA1/IMA2ox* lines showed a more pronounced response, covering a larger root area (Fig. 3A). The enzymatic reduction of ferric chelates is a key step in the strategy I-type of Fe acquisition. In wild-type plants, *in vivo* Fe chelate reduction (FCR) activity was increased by 7-fold when grown on navFe media (Fig. 3B). While FCR activity of *f6’h1* mutant plants was similar to wild-type plants under Fe-sufficient conditions, the mutant showed a much higher (ca. 15-fold) increase in activity when subjected to navFe conditions, possibly caused by the lack of Fe acquired via the secretion of Fe-mobilizing coumarins (Fig. 3B). In both *IMA1ox* and *IMA2ox* lines, FCR activity was significantly increased under Fe-sufficient (ca. 5-fold) and navFe conditions (ca. 2-fold) relative to wild-type plants. Western blot analysis revealed that the Fe^2+^ transporter IRT1 is lowly expressed under Fe-sufficient conditions, with a slight increase in *IMA1ox* and, to a lesser extent, in *IMA2ox* lines when compared to wild-type plants (Fig. 3C). Under navFe conditions, the expression pattern of IRT1 protein resembled that of the *in vivo* FCR activity; highest IRT1 accumulation was observed in*f6’h1* plants and *IMAox* lines (Fig. 3C).

**Figure 3.**
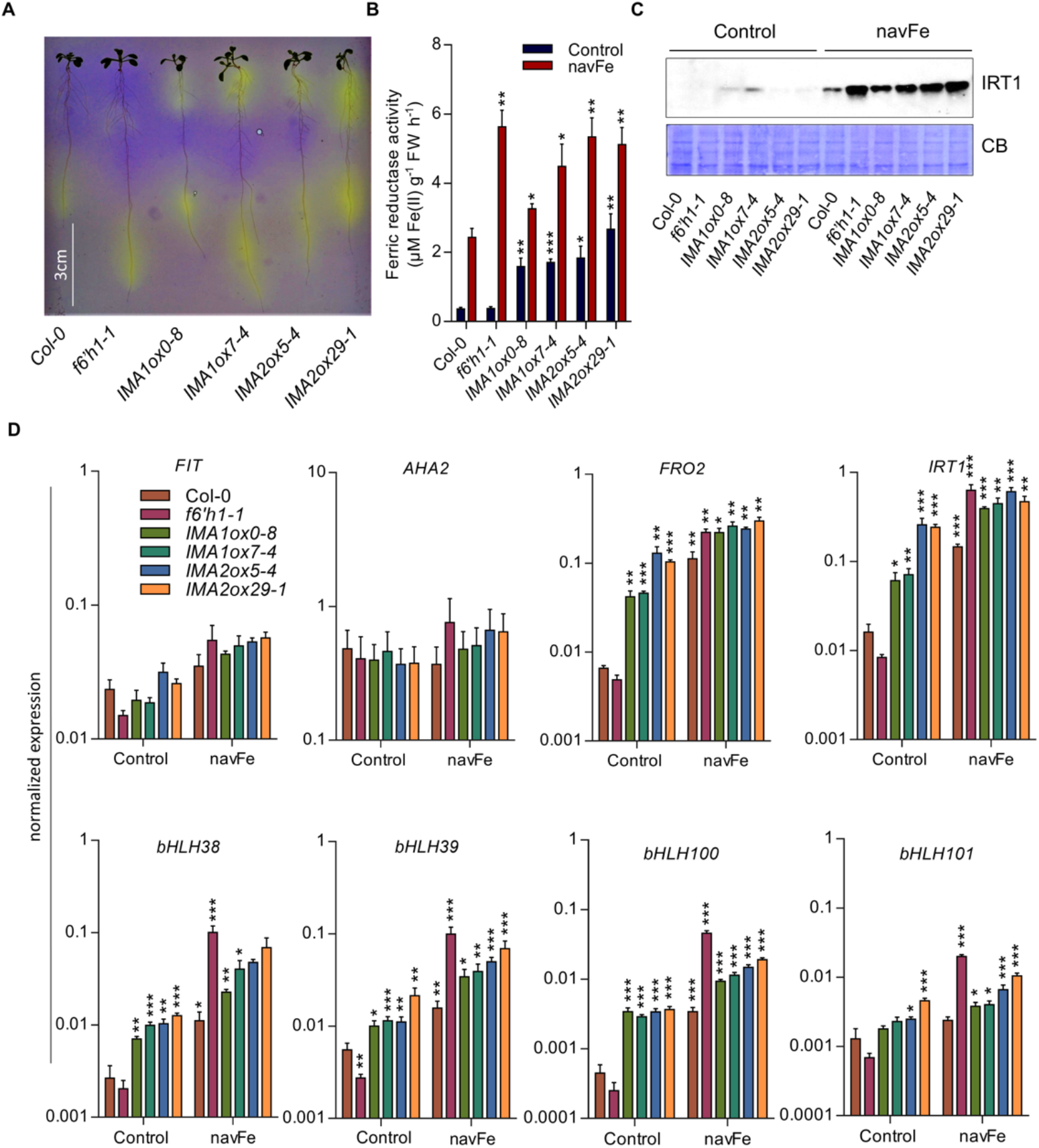
Iron acquisition processes are enhanced in *IMA1* and *IMA2* overexpression lines. A, Rhizosphere acidification by 8-d-old plants grown on ES media after transferring to plates containing the pH indicator bromocresol purple for 24 h. A representative image of three independent experiments is shown. Acidifcation is indicated by yellow color around the roots. B, Ferric reductase activity of 12-d-old plants grown on either ES or navFe media for 3 days after 9 days of pre-cultivation on ES media. C, Expression of IRT1 protein in root samples from 12-d-old plants grown on either ES or navFe media for 3 days after 9 days of pre-cultivation on ES media. Coomassie blue (CB) was used as loading control. Per lane, 10 μg of total protein was loaded. Representative images of three independent experiments are shown. D, Expression of major Fe uptake genes in roots of 12-d-old seedling grown on ES or navFe media for 3 days after 9 days of pre-cultivation on ES media. Gene expression was determined by RT-qPCR using ΔC_T_ method and expression of elongation factor 1 alpha as an internal control. Each bar represents the mean ± SE of four independent experiments. Asterisks indicate significant differences from the wild type in each treatment: *, *P ≤* 0.05; **, *P ≤* 0.01; ***, P ≤ 0.001.

Together, these data show that higher IMA levels activate proton extrusion, ferric reduction, and Fe uptake at different regulatory levels and in conjunction with the Fe status of the plants. The data further indicate that the higher FRO2 activity and IRT1 protein levels of *f6’h1* mutants are insufficient for survival at elevated pH, suggesting that the secretion of Fe-mobilizing coumarins is indispensable to thrive under such conditions. It can further be concluded that the improved growth of *IMAox* lines on media with low Fe availability is not solely caused by induction of the AHA2-FRO2-IRT1 module.

### The Composition of Secreted Coumarins is Dependent on the pH/Fe Regime

The quantities of coumarins in roots and in the media can be non-invasively approximated by detecting autofluorescence as an inherent property of some compounds such as scopoletin, scopolin, and esculin. Under Fe-sufficient conditions, only a faint signal was observed in roots of all genotypes; *IMAox* lines emitted a slightly more intense signal (Supplementary Fig. S4A). When grown on navFe media, a significant increase in autofluorescence was noted for *IMA1/IMA2ox* lines relative to wild-type plants (Fig. 4A,B). In both roots and media of *f6’h1* and *ima8x* mutants, fluorescent coumarins remained below the detection limit under all conditions. Media of *IMA1/IMA2ox* lines not only showed pronounced fluorescence, but also a yellowish color, possibly attributable to the formation of fraxetin-Fe complexes (Rajniak et al, 2018) (Supplementary Fig. S4B).

**Figure 4.**
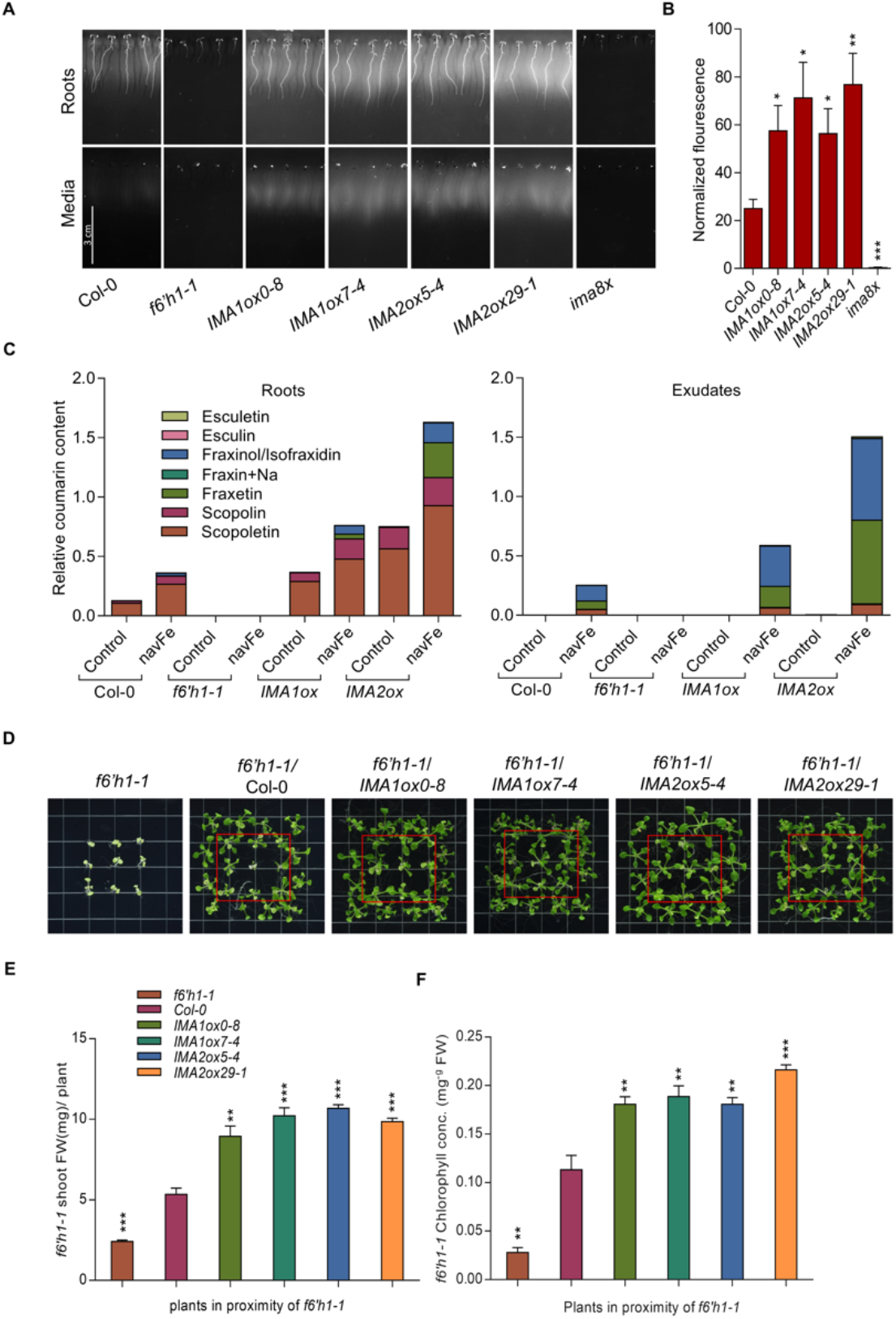
Overexpression of *IMA1* and *IMA2* supports the secretion of Fe-mobilizing coumarins under navFe conditions. A, Accumulation of fluorescent coumarins in roots and media of 7-d-old plants grown on navFe media. Photos were captured before and after removing plants from the media using 365 nm as the excitation wavelength. Representative images of three independent experiments are shown. B, Quantification of root exudate fluorescence. C, Relative abundance of selected coumarins in roots and exudates of plants grown on ES or navFe media. Bars represent the relative content of different coumarin compounds presented as peak area normalized to the number of roots and internal standard peak area. Values are mean relative coumarin content obtained from three independent experiments. D, Rescue of the phenotype of *f6’h1-1* mutant plants (in red boxes) by wild-type, *IMA1ox*, or *IMA2ox* plants grown alongside mutant plants on navFe media. Representative images of four independent experiments are shown. E, Shoot fresh weight of rescued*f6’h1-1* plants. F, Total chlorophyll concentration of leaves from rescued *f6’h1-1* plants. Each bar represents the mean ± SE of four independent experiments. Asterisks indicate significant differences from the wild type in each treatment: *, *P ≤* 0.05; **, *P ≤* 0.01; ***, P ≤ 0.001.Notably, transcript levels of the master regulator *FIT* and the P-type H^+^-ATPase *AHA2* did not significantly differ among the treatments or genotypes under investigation, indicative of a chiefly post-transcriptional regulation of gene activity (Fig. 3D). Instead, expression of *FRO2* and *IRT1* was massively increased in *IMA1*/*IMA2ox* lines under all Fe regimes, with higher transcript levels under navFe conditions. When grown on navFe media, in *f6’h1* plants *FRO2* and *IRT1* mRNA levels were higher than in the wild type and at the level of *IMAox* lines (Fig. 3D). Genes encoding clade Ib bHLH proteins showed a pattern similar to *FRO2* and *IRT1*, with a significant increase under Fe-sufficient conditions in *IMAox* lines except for *bHLH101*. When grown on navFe media, all genes of the clade Ib bHLH clade were significantly higher expressed in *IMAox* lines when compared with the wild type.

Under Fe-sufficient conditions, targeted coumarin analysis identified scopoletin and its glycoside scopolin as the main phenolic compounds in roots. In contrast to the lack of fluorescence under Fe-sufficient conditions, *IMAox* lines showed a marked increase in the level of scopolin and scopoletin, compounds that constitute the major contributors to root fluorescence (Döll et al., 2017) (Fig. 4C). The lack of fluorescence in the roots under Fe-sufficient, slightly acidic condition can be attributed to a pH-dependent shift in the spectra of fluorescent coumarins such as scopoletin, an assumption that was confirmed by UV-Vis spectroscopy (Supplementary Fig. S5A,B). When grown on navFe media, small amounts of fraxinol/isofraxidin were detected in the wild type in addition to scopolin and scopoletin, which account for the largest part of the detected coumarins. In roots of *IMA1/IMA2ox* lines, all identified coumarins were increased relative to the wild type, including the Fe-mobilizing compound fraxetin, which was detectable only in very low quantities in the wild type. *IMA1ox* and *IMA2ox* lines accumulated 28- and 200fold more fraxetin in the roots than the wild type, respectively.

In the media, fraxinol/isofraxidin, and fraxetin were the most prominent compounds in wild-type plants, scopoletin was detected in minor amounts. Similar to what was observed in roots, media of *IMA1*/*IMA2ox* lines contained much higher levels of coumarins than that of wildtype plants, which was particularly pronounced when fraxetin levels are considered. *IMA1ox* and *IMA2ox* lines secreted 3- and 10-fold more fraxetin into the media than wild-type plants, an observation which matches the significantly better performance of these lines on alkaline soil. By contrast, scopoletin levels did not differ much among the lines, except for *f6’h1* mutants which did neither produce nor secrete any detectable amount of the coumarins covered in the analysis (Fig. 4C). The difference in fraxetin secretion between wild-type plants and *IMA1*/*IMA2ox* lines was also observed in experiments aimed at rescuing the mutant phenotype of *f6’h1* plants. Growing *f6’h1* mutant plants alongside Col-0 doubled the shoot fresh weight and increased chlorophyll concentration of the mutant by approximately 4-fold (Fig. 4D-F). Replacing wildtype plants with *IMA1/IMA2ox* lines further improved growth and chlorophyll concentration of *f6’h1* plants, supporting the supposition that the coumarins secreted by *IMA1/IMA2ox* lines are biologically active and qualitatively or quantitatively superior to that of the wild type (Fig. 4D-F).

Together, these data indicate that secretion of Fe-mobilizing compound requires an unidentified Fe deficiency/high pH signal; in contrast to the pronounced induction of proton extrusion, FCR activity, and IRT1 accumulation in the presence of Fe, no coumarins are secreted by *IMA1/IMA2ox* lines under Fe-sufficient conditions.

### The Expression of Genes Involved in Coumarin Biosynthesis is Controlled by pH_e_ and Fe

To further investigate the importance of the Fe/pH regime for the secretion of coumarins, we determined the transcript levels of genes involved in coumarin secretion under Fe-sufficient (pH 5.5) and navFe conditions (pH 7.0). Under Fe-sufficient conditions, the expression of *F6’H1* did not change in lines overexpressing *IMA1*/*IMA2* relative to wild-type plants (Fig. 5A). Growing plants on navFe media increased *F6’H1* mRNA levels significantly with no apparent differences among the genotypes, except for *f6’h1* mutant plants which showed low transcript levels under all conditions. Expression of *S8H,* which mediates the conversion from scopoletin to fraxetin, was increased by overexpression of *IMA* under Fe-sufficient conditions and strongly induced in all lines under navFe conditions, with higher expression in all *IMAox* lines relative to wild-type plants. Conspicuously, an increase in *S8H* transcript levels relative to wild-type plants was also observed in*f6’h1* mutants under navFe but not under Fe-sufficient conditions, possibly due to more depleted Fe levels of the plants under these conditions (Fig. 5A). The strong induction of *S8H* was consistent with higher accumulation of S8H protein in the*f6’h1* mutant and *IMAox* lines (Fig. 5B). While overexpression of *IMA1* and *IMA2* highly increased *CYP82C4* transcript levels under (acidic) Fe-sufficient conditions, such increase was neither observed in wild-type nor in *IMAox* plants when plants were grown under navFe conditions. Instead, in all lines transcription remained at the low levels observed under control conditions (Fig. 5A).

**Figure 5.**
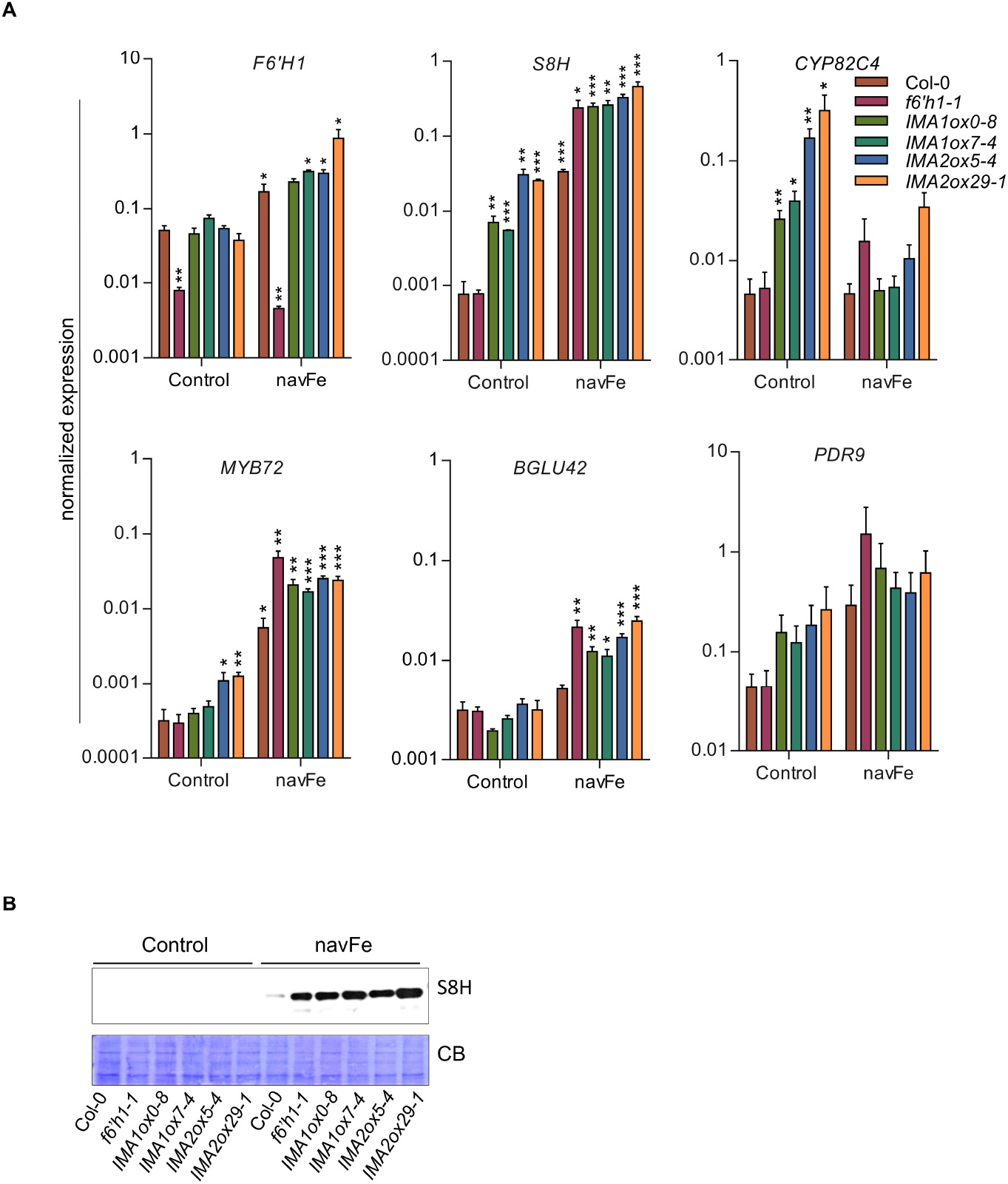
Expression of coumarin biosynthesis-related genes in *IMA1/IMA2ox* lines. RT-qPCR analysis of gene expression in roots of 12-d-old seedling grown on ES or navFe media for 3 days after 9 days preculture on ES media. Gene expression was determined using the ΔC_T_ method with elongation factor 1 alpha as an internal control. Each bar represents the mean ± SE of four independent experiments. Asterisks indicate significant differences from the wild type in each treatment: *, *P* ≤ 0.05; **, *P* ≤ 0.01; ***, P ≤ 0.001. B, Expression of S8H protein in root samples from 12-d-old plants grown on either ES or navFe media for 3 days after 9 days of precultivation on ES media. Coomassie blue (CB) was used as loading control. Per lane, 10 μg of total protein was loaded. Representative images of three independent experiments are shown.

The expression pattern of *MYB72,* a transcription factor that is critical for growth under Fe-limiting, alkaline conditions (Palmer et al., 2013), resembled that of *S8H* except for a lack of induction by overexpression of *IMA1/IMA2* under Fe-sufficient conditions of the former gene (Fig. 5A). Induction of the MYB72-dependent *BETA GLUCOSIDASE42 (BGLU42),* which mediates the deglycosylation of scopoletin and possibly other coumarin glycosides as a necessary step before their secretion into the rhizosphere (Zamioudis et al., 2014), was observed under navFe conditions, but *BGLU42* transcript levels did not differ among the genotypes when grown on ES media. No statistically significant changes were observed for the expression of the coumarin exporter *PLEIOTROPIC DRUG RESISTANCE9 (PDR9),* although a trend towards higher transcript levels was observed under navFe conditions (Fig. 5A).

### Environmental pH Modulates Gene Transcription

To distinguish the effect of pH_e_ from that of Fe availability, we determined the transcript level of genes involved in the secretion of Fe-mobilizing coumarins at various Fe/pH regimes. The activity of Fe in the media is dependent on the stability of the Fe compound, which in turn is strongly affected by pH. To account for this effect, we grew plants on Fe-deficient media (+ferrozine), and media supplemented with various Fe sources, including FeEDTA (unstable at elevated pH), Fe-ethylenediamine-N,N’-bis (FeEDDHA; stable over the pH range under investigation), FeCl_3_ (low solubility particularly at elevated pH), and precipitated Fe(OH)3 (very low solubility over the whole pH range). At pH 5.5, expression of *MYB72* was strongly induced by Fe deficiency and, to a lesser extent, by low Fe availability (Fig. 6A). At pH 7.0, *MYB* transcript levels were much higher under all condition, except for Fe-deficient media where expression appears to reach maximum transcription rates which cannot be further enhanced. The transcription of *BGLU42, F6’H1,* and *COSY* was not much affected by the various growth conditions (Fig. 6A). Instead, *S8H* expression was strongly induced by Fe deficiency at pH 5.5 and by supplementing the media with FeCl_3_ or Fe(OH)_3_ (Fig. 6A). Growth on media with elevated pH increased the message levels of *S8H* under all conditions except for Fe-deficient media, particularly when plants were grown on media supplemented with lowly soluble Fe compounds. Generally, the *S8H* expression pattern resembles that of *MYB72* (Fig. 6A). Western blot analysis mirrored the high induction of *S8H* by Fe deficiency at both acidic and elevated pH and the pH-dependent expression in all treatments, with increasing protein abundance with decreasing stability of the added Fe compounds (Fig. 6B). While expression of *CYP82C4* showed a pattern similar to that of *S8H* at pH 5.5, growth at pH 7.0 strongly downregulated transcript levels in all treatments (Fig. 6A). Although inducibility by Fe deficiency was still observable at neutral pH, expression levels of *CYP82C4* were more than 600-fold lower at pH 7.0 when compared to pH 5.5, suggesting that at elevated pH fraxetin is favored over sideretin production. The expression of *FIT*, *IMA1*, and *IMA2* was not much affected by the Fe/pH regimes under study, except for a strong induction of all three genes by the absence of Fe which was not influenced by pH. Interestingly, expression of *IMA3* showed a different pattern, being strongly repressed by stable Fe compounds at elevated pH (Supplementary Fig. S6).

**Figure 6.**
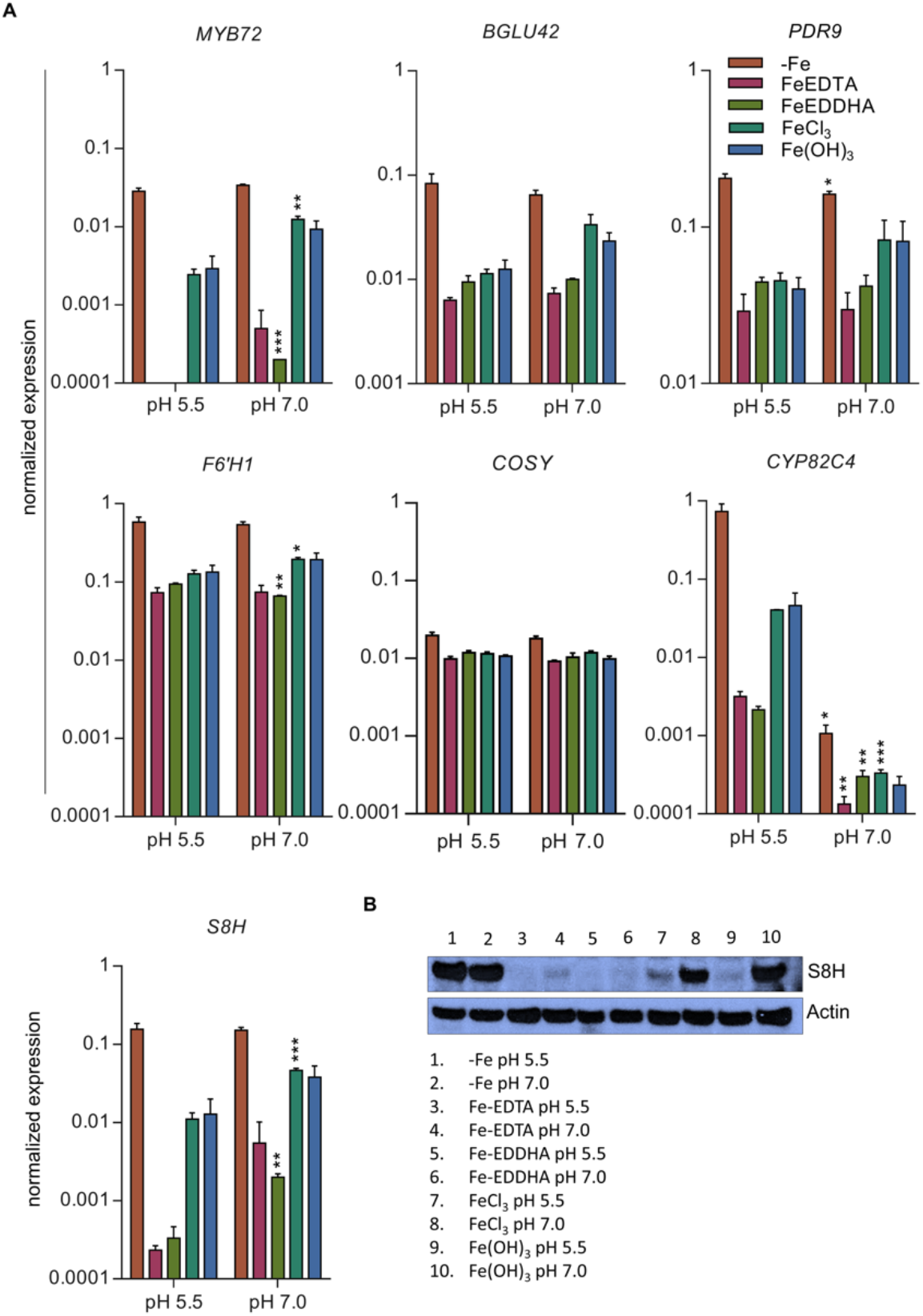
Media pH and Fe availability control the expression of coumarin-related genes. Plants were directly germinated for 14 days on either Fe-deficient media (with 120 μM ferrozine) or media supplemented with either FeEDTA (40 μM), FeEDDHA (40 μM), FeCl_3_ (40 μM), or precipitated Fe(OH)3 (40 μM) at pH 5.5 or pH 7.0. A, Expression of *MYB72, BGLU42, PDR9, F6’H1,COSY, CYP82C4,* and *S8H* in roots. Gene expression was determined using the ΔC_T_ method with elongation factor 1 alpha as an internal control. Each bar represents the mean ± SE of three independent experiments. Asterisks indicate significant differences from pH 5.5 media within the same Fe regime: *, *P ≤* 0.05; **, *P ≤* 0.01; ***, P ≤ 0.001. B, Expression of S8H protein in root samples. Actin was used as loading control. Per lane, 20 μg of total protein were loaded. Representative images of three independent experiments are shown

In conclusion, the results from these experiments suggest that depending on the external pH, *IMA1/IMA2* support the biosynthesis and secretion of different coumarins. Under acidic conditions, sideretin production is favored through upregulation of both *S8H* and *CYP82C4.* At elevated pH and in the absence of available Fe, *S8H* is highly induced while *CYP82C4* expression is strongly repressed, supporting the production of fraxetin which forms more stable complexes with Fe under such conditions.

### IMA1 and IMA2 are Transcriptional Activators

The molecular mechanism by which IMA peptides induce Fe deficiency responses has not been elucidated. To investigate whether IMAs can activate the expression of genes required for the Fe deficiency response, we conducted luciferase reporter assays on selected genes activated by IMA peptides under acidic or alkaline conditions. *CYP82C4* was chosen as a representative key player for IMA-induced genes at acidic pH ranges. For the calcicole branch of the Fe deficiency response, we selected *MYB72* as an upstream regulator for the biosynthesis of catecholic coumarins. Arabidopsis mesophyll protoplasts were transfected with a construct containing either *IMA1* or *IMA2* cDNA under the control of the CaMV 35S promotor *(_pro_35S::IMA1/2)* and plasmid DNA containing the luciferase reporter construct under the *MYB72* or *CYP82C4* promoters *(_pro_MYB72::LUC* and *_pro_CYP82C4::LUC).* IMA firefly luciferase activity was normalized against the activity of Renilla luciferase to correct for transfection efficiency. The luciferase assay revealed that both *IMA* genes strongly stimulated the expression of the two target genes (Fig. 7), suggesting that IMA peptides act as transcriptional regulators. It can also be assumed that the pH signal is perceived downstream of or interferes with the regulation by IMAs.

**Figure 7.**
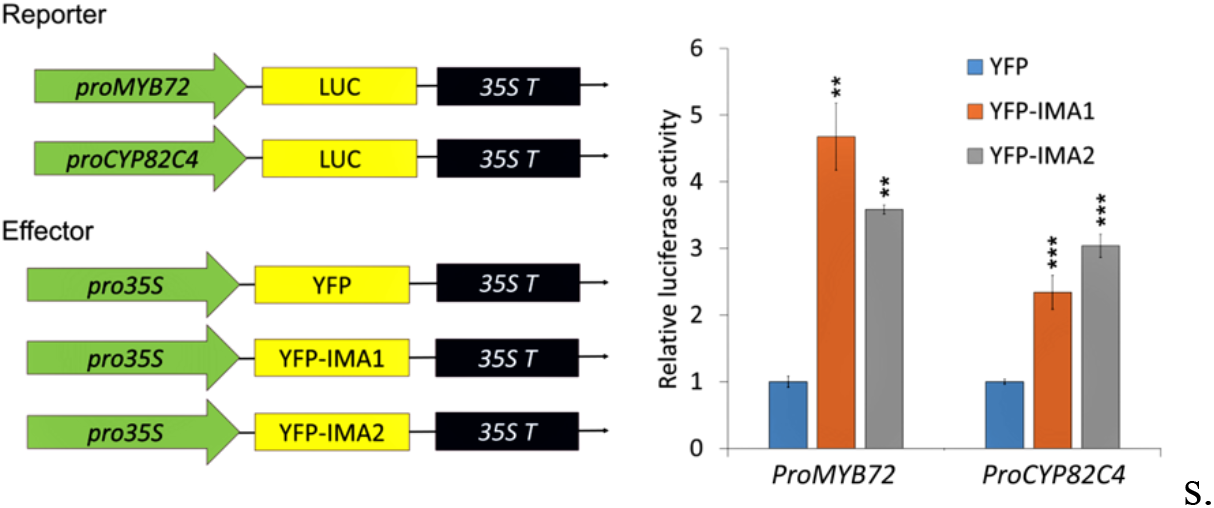
Relative luciferase activity of *MYB72* and *CYP82C4* in Arabidopsis protoplasts cotransfected with *_pro_35S::IMA1, _pro_35S::IMA2,* and *_pro_MYB72::LUC* and *_pro_CPC82C4::LUC*. The data represent SE of three independent experiments. IMA firefly luciferase activity was normalized against the Renilla luciferase activity to correct for transfection efficiency. Asterisks indicate significant differences from the empty YFP: *, *P ≤* 0.05; **, *P ≤* 0.01; ***, P ≤ 0.001.

## DISCUSSION

### *IMA* Overexpression Confers Calcicole Behavior

The Arabidopsis genome harbors eight *IMA* genes, of which only two *(IMA1/FEP3* and *IMA3/FEP1)* have been functionally explored (Grillet et al., 2018; Hirayama et al., 2018). IMA1 and IMA2 share an identical C-terminal consensus motif characteristic of all IMA peptides across angiosperms, exhibit similar expression patterns, and induce Fe acquisition responses in a similar manner and to a comparable extent, suggesting that the two genes are to a large degree genetically redundant. In leaves, *IMA1* and *IMA2* are tightly co-expressed with *IMA4* and genes that have been associated with cellular Fe homeostasis. Notably, in roots, expression of *IMA1/2* is closely associated with *NAS4,* a putative target of MYB72 (Palmer et al., 2013). Homozygous *myb72* and *nas4* mutants display similar phenotypes, featuring lower chlorophyll and decreased Fe levels when compared with the wild type. Overexpression of *NAS4* rescued *myb72* mutants (Palmer et al., 2013), indicating that NAS4 is of particular importance for the adaptation to alkaline conditions with low Fe availability.

The enzymatic reduction of ferric chelates is the rate-limiting step of strategy I-type Fe uptake and determines the efficiency by which Fe is taken up (Grusak et al., 1990). Under Fe-sufficient (and slightly acidic) conditions, elevated IMA levels increase the expression of *FRO2* and *in vivo* FCR activity, which is causative for increased Fe levels in all plant parts including seeds (Grillet et al., 2018). However, activated FCR activity and increased proton extrusion of *f6’h1* plants are insufficient to alleviate the pronounced growth inhibition of the mutant on media with elevated pH, indicating that the secretion of Fe-mobilizing coumarins is indispensable under such conditions. In *IMAox* lines, fraxetin secretion was massively increased under permissive conditions (low Fe status of the plants and elevated pH), an increase that was much more pronounced than that observed for other Fe acquisition responses. Conspicuously, under Fe-sufficient conditions, an IMA-dependent induction was observed for the synthesis of scopolin and scopoletin but not for fraxetin, indicating that IMA-mediated fraxetin production requires both a low Fe status and high pH_e_, thereby prioritizing the secretion of fraxetin over that of the upstream (scopoletin) and downstream products (sideretin). At elevated pH, a concurrent, unweighted induction of all steps of the coumarin pathway may be counterproductive for Fe acquisition, since sideretin secretion will reduce fraxetin levels and thus compromise the mobilization of Fe from recalcitrant pools in soils.

Similar to *IMAox* lines, in wild-type plants the most pronounced accumulation and secretion of fraxetin was observed under the combined influence of elevated pH_e_ and Fe deficiency; no such pattern was observed for scopoletin (Sisó-Terraza et al., 2016). A pH-dependent increase in the secretion of phenolic compounds was also reported for soybean and mung bean plants (Waters et al., 2018; Nair et al., 2020), suggesting that the pH response is conserved in some if not most non-grass species. We show here that in the presence of the pH-stable Fe source FeEDDHA a raise in media pH from 5.5 to 7.0 led to a markedly higher (ca. 6fold on average) expression of *S8H*. Under similar conditions, *CYP82C4* transcript levels were decreased by the same factor. At elevated pH, the Fe deficiency-induced expression of *CYP82C4* was compromised, amplifying the difference in transcript levels between pH 5.5 and pH 7.0 to more than 600-fold. Similar to elevated pH ranges, under acidic conditions an indiscriminate induction of all steps of the coumarin biosynthetic pathway would be disadvantageous for plant fitness; fraxetin synthesis would cause an unfavorable decrease of the phytoalexin scopoletin and decrease the production of sideretin, which would deplete the amount of chelated Fe as a substrate for FRO2-mediated ferric reduction. It should be noted that the expression of both *S8H* and *CYP82C4* is controlled by FIT (Colangelo and Guerinot, 2004) and inducible by Fe deficiency at the transcript and protein level under acidic conditions (Lan et al., 2011; Grillet et al., 2018). This finding implies that *trans-acting* factors can modulate or abolish FIT-dependent induction in response to pH_e_. In contrast to yeast, where ambient pH is perceived by a sensor complex at the cell surface and communicated via a well-described signaling cascade to drive the expression of genes adapting the cell to an alkaline environment (Obara et al., 2014, Peñalva et al., 2014), such pH_e_ signaling has been not yet been described in plants (Raven, 1990; Tsai and Schmidt, 2021). However, changes in media pH were shown to rapidly and thoroughly alter gene expression profiles in Arabidopsis roots, strongly suggesting that plants are capable of monitoring and responding to changes in pH_e_ (Lager et al., 2010; Tsai and Schmidt, 2020). Noteworthily, some of the genes under study did not show sensitivity towards changes in environmental pH, excluding non-specific secondary effects of pH_e_ on gene expression.

### Fe Deficiency Responses Fractionate into a Calcifuge and a Calcicole Branch

The strategy I-type Fe deficiency response constitutes a co-operative entity, which orchestrates Fe acquisition processes at low Fe availability. Our data imply that different branches of the Fe deficiency syndrome are calibrated in a pH_e_-dependent manner, accentuating processes which are most efficient under a given set of edaphic conditions (**Fig. 8**). The balance between the two branches of the Fe deficiency response is tipped by the environmental hydrogen activity, inducing a ‘calcifuge’ suite of processes at acidic pH, and a ‘calcicole’ response at elevated pH.

**Figure 8.**
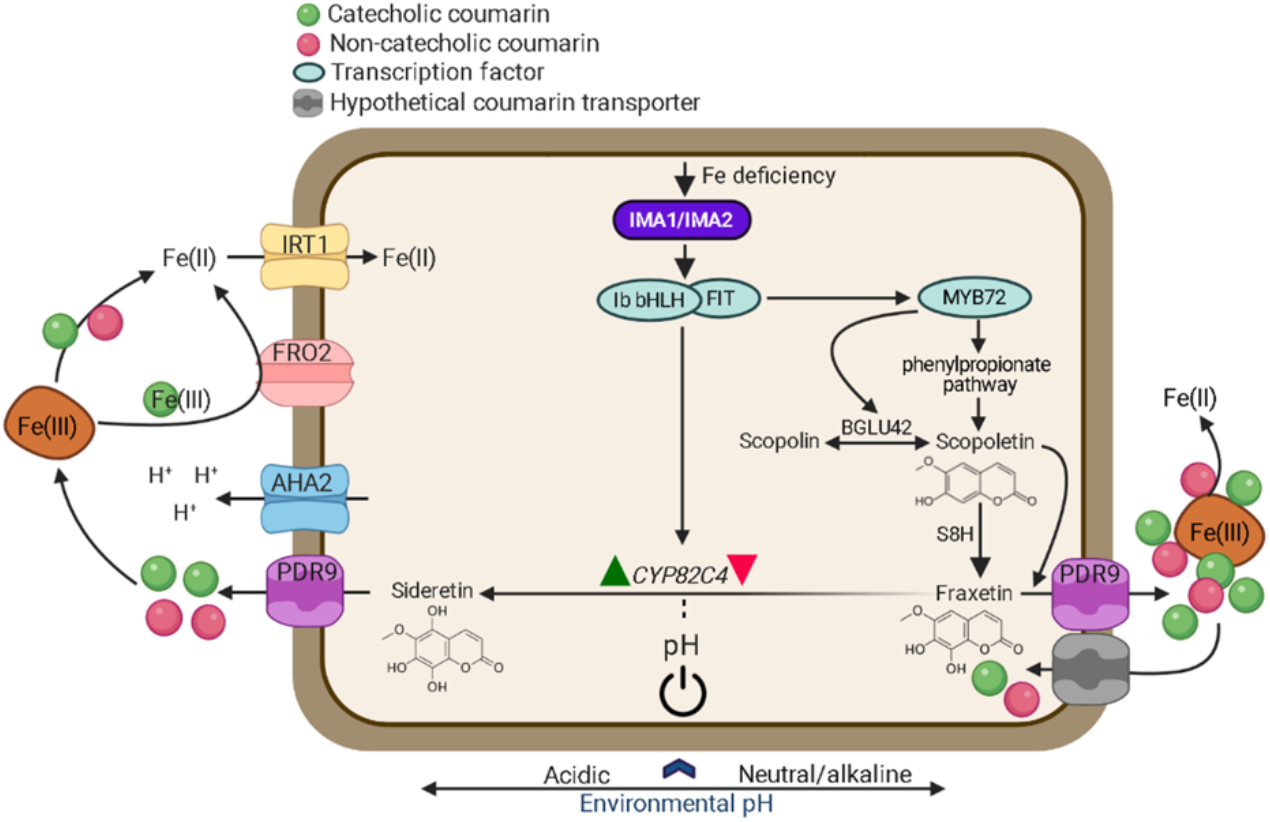
Model for the pH-dependent induction of the calcifuge (left) and calcicole (right) branch of the Fe deficiency response. The expression of *IMA* genes is induced by a low Fe status of the plant, triggering the induction of two distinct suites of responses in correspondence with pH_e_. Environmental pH dictates the biosynthesis of either sideretin or fraxetin by supporting the expression of *CYP82C4* at low pH_e_ and repressing *CYP82C4* transcription at elevated pH_e_ (indicated by green and red arrowheads). At acidic pH, IMAs support the acquisition of Fe from Fe hydroxides by inducing proton extrusion, sideretin secretion, ferric chelate reduction, and uptake of Fe^2+^ via IRT1. The calcicole branch is positively regulated by MYB72. At neutral or alkaline pH ranges, IMAs induce the biosynthesis and secretion of fraxetin via PDR9, and Fe is mobilized via reduction and chelation. The β-glucosidase BGLU42 is prioritized over other BGLUs and required to facilitate the secretion of coumarins by catalyzing their deglycosylation. Re-uptake of fraxetin into the roots as such or as fraxetin-Fe complex is mediated by an as yet unidentified transporter. The two branches are not mutually exclusive, but prioritized by the prevailing pH_e_.

Under acidic conditions, Fe is mainly acquired via enzymatic reduction and taken up through FRO2/IRT1. It can be assumed that CYP82C4 activity and sideretin production is closely associated with the AHA2-FRO2-IRT1 module, a supposition that is corroborated by the extremely close correlation of *IRT1* and *CYP82C4* expression (http://atted.jp), and the identification of CYP82C4 as a potentially interactor of IRT1 (Martín-Barranco et al., 2020). Sideretin exhibit maximal Fe-mobilizing pH capacity at lower pH ranges when compared to fraxetin (Rajanik et al., 2018; Tsai et al., 2018), implying that sideretin secretion is part of the calcifuge branch. A pH-dependent prioritization of Fe acquisition process is of ecological significance, since it profoundly increases the range of hydrogen activities in which Fe can be assimilated from the soil.

The transcription factor *MYB72* was found to be critical for growth on alkaline soil (Palmer et al. 2013). In the present study, we show that *MYB72* expression is highly pH-dependent, exhibiting a pattern similar to *S8H* but contrary to *CYP82C4,* which places MYB72 in the calcicole branch of the response. The β-glucosidase *BGLU42* is a putative target of MYB72 (Palmer et al., 2013), is highly responsive to Fe deficiency (Grillet et al., 2018), and was shown to deglycosylate scopolin (Ahn et al., 2010; Stringlis et al., 2018). Interestingly, under navFe conditions, overexpression of *IMA1/2* significantly increases *BGLU42* expression relative to wild-type plants (**Fig. 5**), while the expression of three other scopoletin-hydolyzing β-glucosidases from the same clade, *BGLU21*, *BGLU22*, and *BGLU23*, was decreased under such conditions (Ahn et al., 2010; Tsai and Schmidt, 2020). Moreover, *BGLU21* and *BGLU22* are downregulated in *IMA1ox* lines under Fe-deficient conditions (Grillet et al., 2018), suggesting that BGLU42 could be involved in the deglycosylation of coumarins including fraxetin at elevated pH_e_. It may thus be assumed that high pH triggers a module of the Fe deficiency response that comprises *MYB72, S8H,* and *BGLU42,* which supports the secretion of fraxetin and confers calcicole behavior (**Fig. 8**). An open question is as to whether secreted coumarins can be transported back into the cell together, either as such or as a complex with Fe. Uptake of the secreted coumarins, in analogy of phytosiderophores released by grasses, would be ecologically advantageous in view of energy conservation and the low efficiency of enzymatic Fe^3+^ reduction at neutral or alkaline pH. Glycosylated coumarins were detected in *f6’h1* plants when grown alongside wild-type plants, or when scopoletin, esculetin, or fraxetin was provided exogenously, suggesting that coumarins can be taken up and stored as glycosides in the vacuole (Robe et al., 2020a). Whether or not the coumarins taken up from the soil solution can be transported into the cell as coumarin-Fe complex by an auxiliary Fe uptake system remains to be elucidated.

### IMAs Act as Transcriptional Co-Activators

In Arabidopsis, all Fe deficiency responses are regulated by a complex cascade of regulatory processes that orchestrate a subtle balance of promotive and repressive *trans*-acting factors in concert with post-translational control of central regulatory nodes (Rodrígues-Celma et al., 2019; Schwarz and Bauer, 2020; Spielmann and Vert, 2020; Gao and Dubos, 2020). Overexpression of *IMA1-IMA3* was shown to induce large parts of the Fe deficiency response in Arabidopsis (Grillet et al., 2018; Hirayama et al., 2018; this study), but the exact position of IMA peptides in the regulatory network governing cellular Fe homeostasis has not yet been defined. We show here that IMA1/IMA2 can activate the transcription of key genes of both the calcifuge and calcicole branch of the strategy I-type response, suggesting that IMAs act downstream of or in combination with known players in Fe homeostatic control. This does not, however, rule out a regulatory function of IMA peptides at more upstream positions. Stabilization of upstream regulatory proteins by, for instance, a delayed or compromised degradation or a function as co-transcriptional activators may be causative for the observed induction of Fe uptake by IMAs.

While attempts to identify DNA-binding properties of IMAs have been unsuccessful so far, the current data support a scenario in which IMAs are translated and control transcript levels of both regulatory factors such as MYB72 and downstream actors such as CYP82C4. Our data also suggest that pH_e_ can interfere with and compromise IMA action, possibly via a *trans-acting* factors which modulate the expression of pH-sensitive genes. In an alternative, not mutually exclusive scenario, subtle changes in internal pH induced by larger alterations in pH_e_, may trigger a signal transduction pathway that relays information on pH_e_ from the apoplast to the nucleus. A candidate for an internal pH sensor is phosphatidic acid (PA), which was shown to integrate cellular pH dynamics with stress responses via protonation and de-protonation of PA, thereby changing its target binding in signal transduction cascades (Li et al., 2019).

## CONCLUSIONS

In summary, we show here that high IMA1/IMA2 levels partly overrule the downregulation of Fe uptake at sufficient internal Fe levels and induce rhizosphere acidification, Fe^3+^ reduction, Fe^2+^ uptake, and the biosynthesis and secretion of scopoletin by acting as transcriptional activators. A complex, pH-dependent regulation was observed for the secretion of Fe-mobilizing catecholic coumarins, prioritizing the production of fraxetin over sideretin at elevated pH. Notably, at elevated pH, *f6’h1* (defective in secreting catecholic coumarins) and *ima8x* mutants (defective in all Fe deficiency responses) showed similar phenotypes, indicating that a functional AHA2-FRO2-IRT1 module is insufficient to enable growth under such conditions.

Root-secreted coumarins not only act directly on Fe uptake via mobilization of recalcitrant Fe pools, but also indirectly by retracting pathogens and attracting beneficial bacteria, which in turn improve the access to mobile Fe in soils (Stringlis et al., 2019; Voges et al., 2019; Harbort et al., 2020). While such beneficial effects on plant growth were shown for scopoletin and fraxetin, sideretin does not appear to play such a role (Harbort et al., 2020). It may be concluded that the dual function of fraxetin in Fe acquisition is critical for conferring fitness at elevated pH. Manipulating the expression of IMA/FEP peptides constitutes a promising approach to generate germplasm adapted to low Fe availability. Ectopic expression of *IMAs* results in an increase in Fe in both strategy I and strategy II crops such as rice and tomato with increased Fe content in grains and fruits, respectively (Grillet et al., 2018; Kobayashi et al., 2020), making IMAs a promising candidate for improving Fe acquisition of virtually all crops. In particular, crops grown on calcareous soils may profit from improved growth and nutritional quality of edible plant parts.

## MATERIALS AND METHODS

### Plant Materials and Growth Conditions

Seeds of Arabidopsis *(Arabidopsis thaliana* (L.) Heynh) accession Columbia (Col-0) and *f6’h1-1* (Salk_132418C) T-DNA insertion mutants were obtained from the Arabidopsis Biological Resource Center (Ohio State University). The lines carrying *CaMV _pro_35S::IMA1,* and *ima8x* octuple mutants have been described previously (Grillet et al., 2018). *Agrobacterium tumefaciens-mediated* transformation (Clough and Bent, 1998) was used to transform wild-type plants with pH2GW7 (obtained from the VIB-UGent Center for Plant Systems Biology) containing CaMV 35S promoter-driven *IMA2* CDS (using the primers listed in Supplemental Table S1), yielding *IMA2* overexpression lines. Overexpression in transgenic plants was confirmed by RT-qPCR. Plants were grown in a growth chamber on solid nutrient media as described by Estelle and Somerville (1987) (ES media), composed of 5 mM KNO_3_, 2 mM MgSO_4_, 2 mM Ca(NO_3_)_2_, 2.5 mM KH_2_PO_4_, 70 μmM H_3_BO_3_, 14 μM MnCl_2_, 1 μM ZnSO_4_, 0.5 μM CuSO_4_, 0.01 μM CoCl_2_, 0.2 μM Na_2_MoO_4_, 1% (w/v) MES, and 43.8 mM sucrose, solidified with 0.4% Gelrite pure (Kelco). Unless stated otherwise, media were supplemented with 50 μM FeEDTA or 400 μM Fe-EDTA (Fe excess; Fe++). The pH was adjusted to 5.5. Media with different Fe sources were prepared by replacing FeEDTA with 40 μM of either FeEDDHA, FeCl_3_, or Fe(OH)_3_. For Fe-deficient media, 120 μM ferrozine were added to the media. For preparing non-available Fe (navFe) media, 10 μM FeCl_3_ were added as an Fe source and the pH was adjusted to 7.0 with KOH. For all media adjusted to pH 7.0, MOPS (1 g/L) was used as buffering agent instead of MES. Seeds were surface-sterilized with 30% (v/v) commercial bleach containing 6% NaClO and 70% (v/v) ethanol for 5 min, followed by five rinses with sterile MQ water, and stratified for 2 d in the dark at 4°C before transferring to a growth chamber. Plants were grown at 21°C under continuous illumination (50 μmol m^2^s^-1^). Soil of variable alkalinity was prepared by mixing 0.5% or 1% CaO (w/w) with peat-based soil.

### RT-qPCR

Unless stated otherwise, 9-d-old plants were transferred from ES media to either fresh ES or navFe media for 3 days. At the end of the experimental period, samples were immediately frozen in liquid nitrogen and stored at −80°C. Total RNA was extracted using the RNeasy Mini Kit (Qiagen) and treated with DNase using TURBO DNA-free kit (Ambion). cDNA was synthesized using DNA-free RNA with oligo-dT (20) primers and SuperScript III First-Strand Synthesis System for RT-qPCR (Invitrogen). After incubation at 50°C for 1 h and 70°C for 15 min, 1 μL of RNase H was added and samples were incubated for 20 min at 37°C. The first-strand cDNA was used as a PCR template in a 10 μL reaction system using the SYBR Green PCR Master Mix (Applied Biosystems) with programs recommended by the manufacturer in an AB QuantStudio Real-Time PCR System (Applied Biosystems). Three independent replicates were performed for each sample. The ΔC_T_ method was used to determine the relative amount of gene expression, with the expression of elongation factor 1 alpha (EF1α; At5g60390) as an internal control. Primers used for RT-qPCR are listed in Supplemental Table 1.

### Biomass and Chlorophyll Concentration

For biomass determination, whole shoots of 12- or 14-d-old seedlings grown on ES or navFe media were harvested and chlorophyll was extracted with 80% acetone. Total chlorophyll was calculated from absorbance measured at 645 and 663 nm using a protocol from Wellburn (1994).

### Fe Concentration

Total Fe concentration in shoots and roots was determined using a protocol described by Schmidt (1996). Shoot and root samples (20-30 seedlings per sample) were oven-dried at 55°C for 2 d and digested with 65% (v/v) HNO_3_ at 100°C for 6 h, followed by 30% (v/v) H_2_O_2_ at 56°C for 2 h. Digested samples were mixed in assay solution containing 1 mM bathophenanthroline disulfonate (BPDS), 0.6 M sodium acetate, and 0.48 M hydroxylamine hydrochloride. The concentration of the resulting Fe^2+^-BPDS_3_ complex was measured at A535 using a PowerWave XS2 microplate spectrophotometer (BioTek). The Fe concentration in different samples was calculated by plotting OD values against a standard curve made with FeCl_3_.

### Perls Staining for Fe(III)

Two-week-old Arabidopsis seedlings were vacuum infiltrated with Perls solution (2% HCl and 2% K_4_Fe(CN)_6_) for 30 min as described by Roschzttardtz et al. (2009). Samples were then rinsed three times with distilled water. Following staining, chlorophyll was removed from leaf samples with 80% acetone. Samples were rinsed thrice with distilled water and micrographs were taken with a Zeiss Lumar V12 Stereo microscope.

### Rhizosphere Acidification

For visualizing rhizosphere acidification, seedlings were germinated vertically on ES media with 0.5 g/L MES (instead of 1.0 g/L used in ES media) for 7 d and then carefully transferred to 1% BACTO^TM^ agar media containing 0.006% Bromocresol Purple and 0.2 mM CaSO_4_ for 24 h. The pH was adjusted to 6.1 with NaOH. Pictures were taken under white light at the end of the experimental period.

### Ferric Chelate Reduction Activity

*In vivo* root ferric chelate reduction (FCR) activity was measured on 12-d-old seedlings after 3 d of growth on either Fe-replete or Fe-deplete media as described previously (Grillet et al., 2014). FCR activity was determined individually on roots of 5 to 10 intact seedlings. Plants were incubated in the dark for 30 mins with mild shaking in 2 mL assay solution consisting of 100 μM FeEDTA and 300 μM BPDS in 10 mM MES at pH 5.5. Spectrophotometric absorbance at 535 nm was recorded with a PowerWave XS2 plate reader (BioTek Instruments) to determine Fe^2+^-BPDS3 concentrations using a standard curve. Experiments were conducted at least three times independently.

### Protein Extraction and Western Blot Analysis

Total protein from liquid-nitrogen-ground root tissues was extracted as described previously (Tsai et al., 2018). Equal amounts of total protein were loaded and separated on Nu-PAGE 4-12% Bis-Tris gels (Invitrogen). Proteins were immune-detected using anti-IRT1 (Agrisera; AS11 1780) or anti-S8H (Tsai et al., 2018) as primary antibodies followed by detection with secondary antibodies (Agrisera; AS09 602, and GE Healthcare; A934-100UL, respectively). The chemiluminescence signal detection was performed using Pierce ECL Western Blotting Substrate (Thermo Fisher Scientific) on X-ray film. Monoclonal anti-actin (plant) antibody detected by the secondary antibody (GE Healthcare; NA931-1ML) was used as a loading control (Sigma-Aldrich; A0480).

### Detection of Fluorescent Compounds in Roots and Media

The accumulation of fluorescent compounds in the roots and secreted into the media was visualized with a Bio Spectrum 600 imaging system (UVP) at 365 nm excitation wavelength, SYBR Gold 485 to 655nm as emission filter, and exposure time set to 9 s as described previously (Tsai et al., 2018). The fluorescent intensity was quantified using the ImageJ tool. To record the UV-Vis spectra, a Varian Carry 5000 spectrophotometer equipped with a 1 cm quartz cuvette was used.

### Coumarin Extraction from Roots and Media

Coumarins were extracted following a protocol adapted from Tsai et al. (2018). For each sample, 18-20 frozen roots in liquid nitrogen were finely ground using glass beads in a Tissue Lyzer II (Qiagen). Subsequently, extraction was performed by adding 500 μL of 80% (v/v) methanol containing 0.5 ppm 4-methylumbelliferone as an internal standard into the finely ground samples. Samples were vigorously vortexed for 15 min. Following centrifugation at 13,200 rpm for 5 min, supernatants were collected in light-protected Eppendorf tubes. The pellets were subjected to re-extraction to obtain a final pooled volume of 1 mL. For coumarin extraction from growth media, roots were removed from the plates and the media were oven-dried at 50°C for 2 d, and subsequently incubated in 10 mL of 80% (v/v) methanol containing 0.1 ppm 4-methylumbelliferone for 30 min on a shaker. The extract was collected without disturbing the media debris. The extraction process was repeated twice. The samples were centrifuged at 13,200 rpm for 10 min and supernatants were used for targeted UPLC analysis.

### Targeted Coumarin Analysis

Targeted coumarin profiling and data analysis was done as previously described (Tsai et al., 2018). UPLC-QTOF-MS analysis was performed on an Acquity UPLC system (Waters) and a SYNAPT G2 high-definition mass spectrometry system (Waters) with an electrospray ionization interface, ion mobility, and time-of-flight system. Spectra were collected in the positive ionization (ES+) mode. To obtain the relative coumarin levels, the metabolite peak area was normalized to the internal standard peak area and the number of roots.

### Dual Luciferase Reporter Assay

Arabidopsis protoplasts were transfected with _pro_35S::YFP, _pro_35S::RenillaLUC and different pLUC constructs *(_pro_MYB72::LUC* and *_pro_CYP82C4::LUC)* using the pHGWL7 vector. The transfected protoplasts were incubated at 23°C for 18 h in the dark, and luciferase activities were measured using the Dual-Luciferase Reporter Assay System (Promega) with the help of Power Wave XS (Bio-TEK). The Renilla luciferase activity was used to normalize the Firefly luciferase activity. Primers for cloning promoters and IMA1/IMA2 CDS are listed in Supplemental Table S1.

## ACKNOWLEDGMENTS

The authors would like to thank Tushar S. Jadhav (Institute of Chemistry, Academia Sinica, Taipei) for insightful discussion on UV-Vis spectra analysis, Dr. Louis Grillet (Department of Agricultural Chemistry, National Taiwan University, Taipei) for generating the *IMA2* overexpression lines without tag constructs, and Dr. Kuo-Chen Yeh (Agricultural Biotechnology Research Center; ABRC, Academia Sinica) for providing the pHGWL7 vector. We thank the Plant Tech Core Facility at ABRC for protoplast isolation and transformation, the Small Molecule Metabolomics core facility, sponsored by the Institute of Plant and Microbial Biology (IPMB), and the Scientific Instrument Center, Academia Sinica, for providing technical assistance, useful discussion, and support with data acquisition and analysis of Synapt HDMS experiments. We also acknowledge the use of the QuantStudio 12K Flex Real-Time PCR System at the Genomic Technology Core Facility at IPMB. The authors are most grateful for the continued support provided by Dr. Wendar Lin from the Bioinformatics Core Facility at IPMB, Academia Sinica.

## Accession Numbers

Arabidopsis Genome Initiative locus identifiers for the genes mentioned in this article are as follows: IMA1, At1g47400; IMA2, At1g47395; IMA3, At2g30766; IMA4, AT1G07367; IRP3, At2g14247; IRP4, At1g13609; IRP5, At3g56360; IRP6, At5g05250; AHA2, At4g30190; S8H, At3g12900; F6’H1, At3g13610; FIT, At2g28160; FRO2, At1g01580; IRT1, At4g19690; MYB72, At1g56160; CYP82C4, At4g31940; bHLH38, At3g56970, bHLH39, At3g56980; bHLH100, At2g41240; bHLH101, At5g04150; EF1α, At5g60390; BGLU42, At5g36890; PDR9, At3g53480; COSY, At1g28680; NAS4, At1g56430; bHLH18, At2g22750; bHLH19, At2g22760; bHLH20, At2g22770; bHLH25, At4g37850; URI, At3g19860; ILR3, At5g54680, BTSL1, At1g74770; BTSL2, At1g18910.

## Supplemental Data

The following supplemental materials are available.

**Supplemental Table S1.** List of primers used in this study.

**Supplemental Figure S1.** Expression patterns of *IMA1* and *IMA2.*

**Supplemental Figure S2.** Perls staining of Col-0, *f6’h1-1, IMA1ox* and *IMA2ox* seedlings.

**Supplemental Figure S3.** Phenotypes and Fe concentrations of Col-0, *f6’h1-1, IMA1ox* and *IMA2ox* seedlings.

**Supplemental Figure S4.** Autofluorescence of seedlings and media.

**Supplemental Figure S5.** pH dependence of scopoletin fluorescence.

**Supplemental Figure S6.** RT-qPCR analysis of key genes involved in Fe homeostasis.

